# Complementary Models of Cardiometabolic Stress Reveal Conserved Molecular Programs Driving Cardiac Remodeling

**DOI:** 10.64898/2026.08.28.747839

**Authors:** Maaria Saeed, Hyun-Jung Jung, Bo Ryung Lee, Swapna Patil, Ripon Sarkar, Connor Lantz, Mi Jeong Heo, Alexandra Serrato, Yu A An, Kang Ho Kim, Matthew DeBerge

**Author notes:** Co-first authors (Maaria Saeed, Hyun-Jung Jung, Bo Ryung Lee).

## Abstract

**Background:** Cardiometabolic diseases frequently involve concurrent cardiovascular and hepatic dysfunction, yet the conserved molecular mechanisms underlying these systemic responses remain poorly defined.

**Objectives:** To identify conserved molecular responses across complementary manifestations of cardiometabolic stress and determine whether integrated multi-organ analyses reveal therapeutically actionable targets for heart failure.

**Methods:** Cardiac functional phenotyping, hepatic injury profiling, and bulk RNA sequencing were performed across three complementary mouse models representing distinct manifestations of cardiometabolic stress: high-fat diet plus L-NAME (HFD+LN)-induced heart failure with preserved ejection fraction (HFpEF; cardiovascular disease), Western diet (WD)-induced obesity (systemic metabolic stress), and choline-deficient, L-amino acid-defined, high-fat diet (CDAHFD)-induced steatotic liver disease (hepatic metabolic stress). Comparative transcriptomic analyses distinguished organ-specific responses from conserved molecular signatures.

**Results:** Each model produced distinct systemic, hepatic, and cardiac phenotypes accompanied by divergent transcriptional responses within individual organs. Cross-model and cross-organ integration identified a limited set of conserved molecular responses to cardiometabolic stress, with *Serpine1*, encoding plasminogen activator inhibitor-1 (PAI-1), emerging as a highly conserved candidate that exhibited preferential induction in the heart. Pharmacologic inhibition of PAI-1 significantly improved cardiac function and attenuated adverse remodeling in established HFpEF, whereas hepatic pathology was comparatively less affected, indicating differential organ-specific dependence on this pathway.

**Conclusions:** Integrated analyses across complementary manifestations of cardiometabolic stress identified conserved molecular signatures that transcend individual disease models and organs. These findings establish a comparative framework for discovering cardiovascular therapeutic targets and identify PAI-1 as a promising mediator of cardiac remodeling in HFpEF.

## INTRODUCTION

Heart failure affects more than 60 million people worldwide and continues to increase in parallel with the global epidemics of obesity, type 2 diabetes, and metabolic syndrome (1). Among heart failure subtypes, heart failure with preserved ejection fraction (HFpEF) represents the fastest growing phenotype and is characterized by limited disease-modifying therapies and persistently high morbidity and mortality (2). Metabolic dysfunction similarly drives chronic liver disease, with metabolic dysfunction-associated steatotic liver disease (MASLD) emerging as one of the most prevalent hepatic disorders worldwide (3). Clinical studies consistently demonstrate that MASLD severity independently predicts incident heart failure, hospitalization, and cardiovascular mortality, while patients with established heart failure exhibit a disproportionately higher prevalence of hepatic steatosis, fibrosis, and biochemical liver injury (4–6). Together, these observations suggest that cardiovascular and hepatic disease represent parallel manifestations of systemic cardiometabolic dysfunction rather than isolated organ-specific disorders.

Experimental models have identified numerous mechanisms contributing to cardiometabolic disease, including chronic inflammation, endothelial dysfunction, fibrosis, metabolic inflexibility, oxidative stress, and innate immune activation (7–10). Although these processes have largely been investigated within individual organs, many are shared across tissues exposed to metabolic stress (11). Consequently, distinguishing conserved pathogenic mechanisms from tissue-specific adaptations remains a major challenge. Most studies have focused on individual disease models or isolated signaling pathways, limiting the ability to identify molecular responses that generalize across diverse cardiometabolic conditions. Comparative analyses spanning complementary manifestations of cardiometabolic stress therefore provide a unique opportunity to distinguish shared responses from organ-specific adaptations, identify conserved pathways that represent fundamental drivers of disease, and accelerate the development of novel therapeutic strategies for cardiovascular disease.

In the present study, we integrated systemic phenotyping, cardiac and hepatic functional assessment, and multi-organ transcriptomic profiling across three complementary preclinical models representing distinct manifestations of cardiometabolic stress: high-fat diet plus L-NAME (HFD+LN) as a HFpEF cardiovascular disease model (12), Western diet (WD)-induced obesity as a model of systemic metabolic stress (13), and choline-deficient, L-amino acid-defined, high-fat diet (CDAHFD)-induced steatotic liver disease as a model of hepatic metabolic stress (14). As discussed below, we define organ-specific and conserved molecular responses to cardiometabolic stress and demonstrate that cross-model, multi-organ integration uncovers therapeutically actionable pathways that are not apparent from individual disease models alone. These findings establish a comparative framework for identifying cardiovascular therapeutic targets that transcend traditional organ- and model-specific approaches.

## RESULTS

### Complementary models recapitulate distinct systemic manifestations of cardiometabolic stress

To establish complementary manifestations of cardiometabolic stress, we compared mice exposed to HFD+LN as a commonly used model of HFpEF (7,12), WD as a model of systemic metabolic stress, and CDAHFD as a model of steatotic liver disease with chow-fed controls (Figure 1A). Body weight increased in HFD+LN and WD treated mice, whereas CDAHFD did not produce comparable weight gain (Figure 1B). Consistent with L-NAME-induced hypertension (15), systolic and diastolic blood pressure were selectively increased in HFD+LN treated mice but remained unchanged following WD or CDAHFD treatment (Figure 1C). Circulating lipid profiles also differed among models. Serum cholesterol was increased in HFD+LN and WD treated mice but not following CDAHFD, while triglyceride concentrations exhibited a distinct model-dependent pattern (Figure 1D). Targeted profiling of circulating free fatty acids further demonstrated differential abundance of individual lipid species across HFD+LN, WD, and CDAHFD treatment (Figure 1E). These findings establish physiologically distinct manifestations of cardiometabolic stress despite the high dietary lipid exposure shared across models.

**Figure 1.**
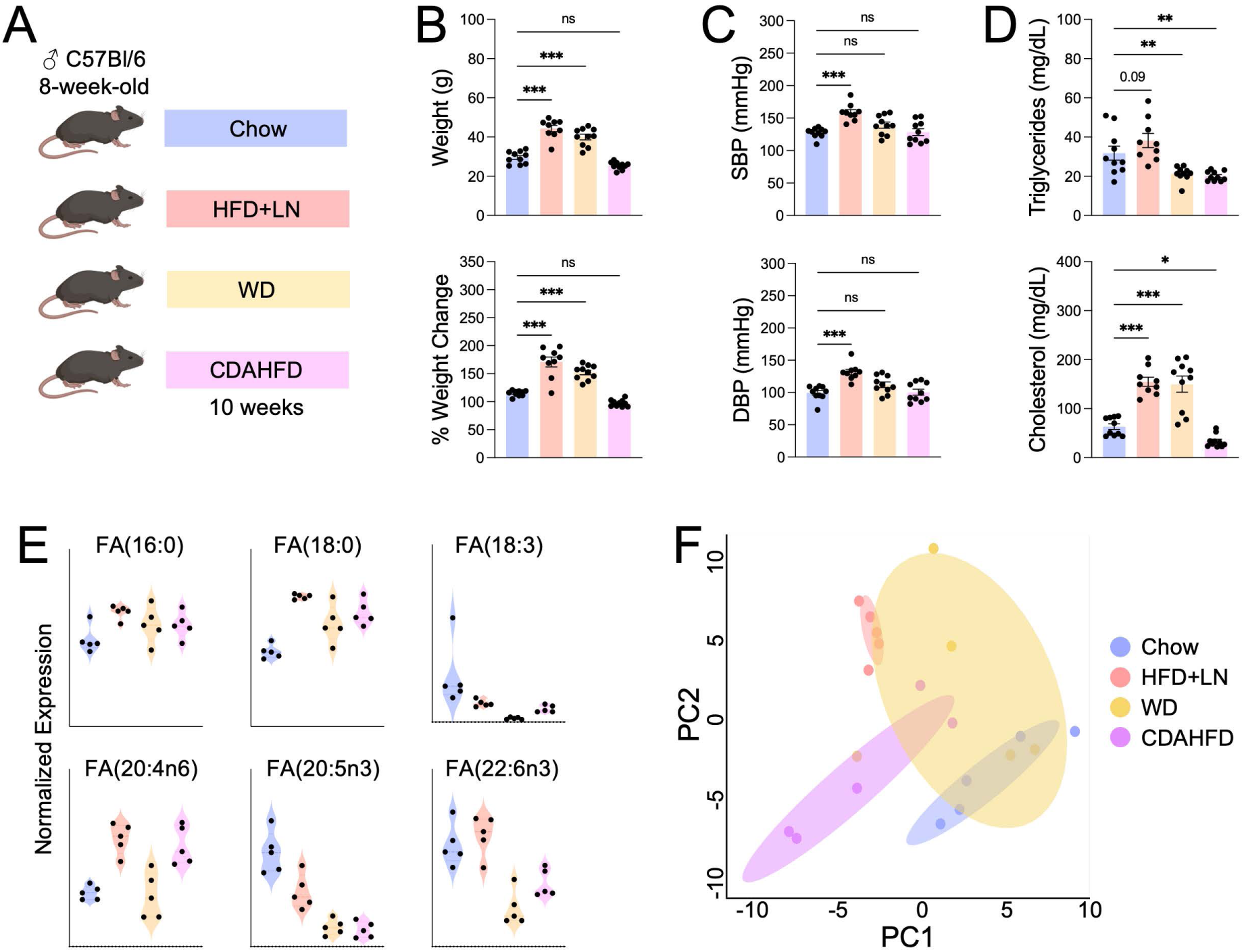
Complementary models of cardiometabolic stress produce distinct systemic metabolic phenotypes. **A** Experimental schematic showing 8-week-old male C57BL/6 mice fed chow, high-fat diet plus L-NAME (HFD+LN), Western diet (WD), or choline-deficient, L-amino acid-defined, high-fat diet (CDAHFD) for 10 weeks. **B** Terminal body weight and percent weight change. **C** Systolic (SBP) and diastolic (DBP) blood pressure. **D** Serum triglyceride and total cholesterol concentrations. **E** Serum free fatty acid concentrations. **F** Principal component (PC) analysis of serum metabolomic profiles. Data are presented as mean ± SEM. *n =* 9-10 mice/group pooled from two or more independent experiments for panels B-D and *n* = 5 mice/group from a single experiment for panels E and F. *\*P <* 0.05, *\*\*P <* 0.01, ***P < 0.001 by one-way ANOVA followed by Tukey’s test.

To determine whether these phenotypic differences were reflected in the circulating metabolic environment, we performed untargeted serum metabolomics. Principal component analysis demonstrated separation among experimental groups (Figure 1F), while differential metabolite analysis identified model-dependent changes spanning amino acid, fatty acid and lipid, nucleotide, nucleotide-sugar, pentose phosphate, and tricarboxylic acid cycle metabolism (Supplemental Figure 1). Notably, HFD+LN treatment increased circulating succinate, consistent with human metabolomic studies demonstrating elevated plasma succinate and other tricarboxylic acid cycle intermediates in patients with HFpEF (16). In contrast, CDAHFD produced broad alterations in amino acid, lipid, and intermediary metabolism, metabolic classes similarly disrupted in human MASLD and MASH (17). Together, these findings establish distinct systemic and circulating metabolic environments across complementary models of cardiometabolic stress, providing a framework to determine how these divergent disease states impact individual target organs.

### Cardiometabolic stress induces shared hepatic injury but distinct inflammatory and fibrotic remodeling

Given these distinct systemic phenotypes, we asked whether complementary forms of cardiometabolic stress produced shared or divergent hepatic responses. Despite differences in overall disease phenotype, HFD+LN, WD, and CDAHFD each produced evidence of liver injury, including increased serum aspartate aminotransferase (AST), hepatic steatosis, and triglyceride accumulation (Figures 2A-D). These changes were most pronounced following CDAHFD treatment, which produced extensive macrovesicular steatosis and disruption of normal hepatic architecture. Consistent with altered lipid handling, *Cd36* expression was increased across disease conditions, with broader changes in fatty acid metabolic genes predominantly observed following HFD+LN and WD treatment (Figure 2E). Notably, *Cyp2c29* was reduced across all three models (Supplemental Figure 2), suggesting disruption of endobiotic and xenobiotic metabolism even in the absence of advanced liver disease. In contrast, robust inflammatory and fibrotic gene expression, together with activation of proinflammatory interferon-associated signaling, was largely restricted to CDAHFD (Figure 2E; Supplemental Figure 2). Thus, while hepatic injury, steatosis, and altered lipid metabolism were shared across cardiometabolic stress models, progression to inflammatory and fibrotic remodeling was predominantly observed with CDAHFD, consistent with the liver-predominant phenotype of this model.

**Figure 2.**
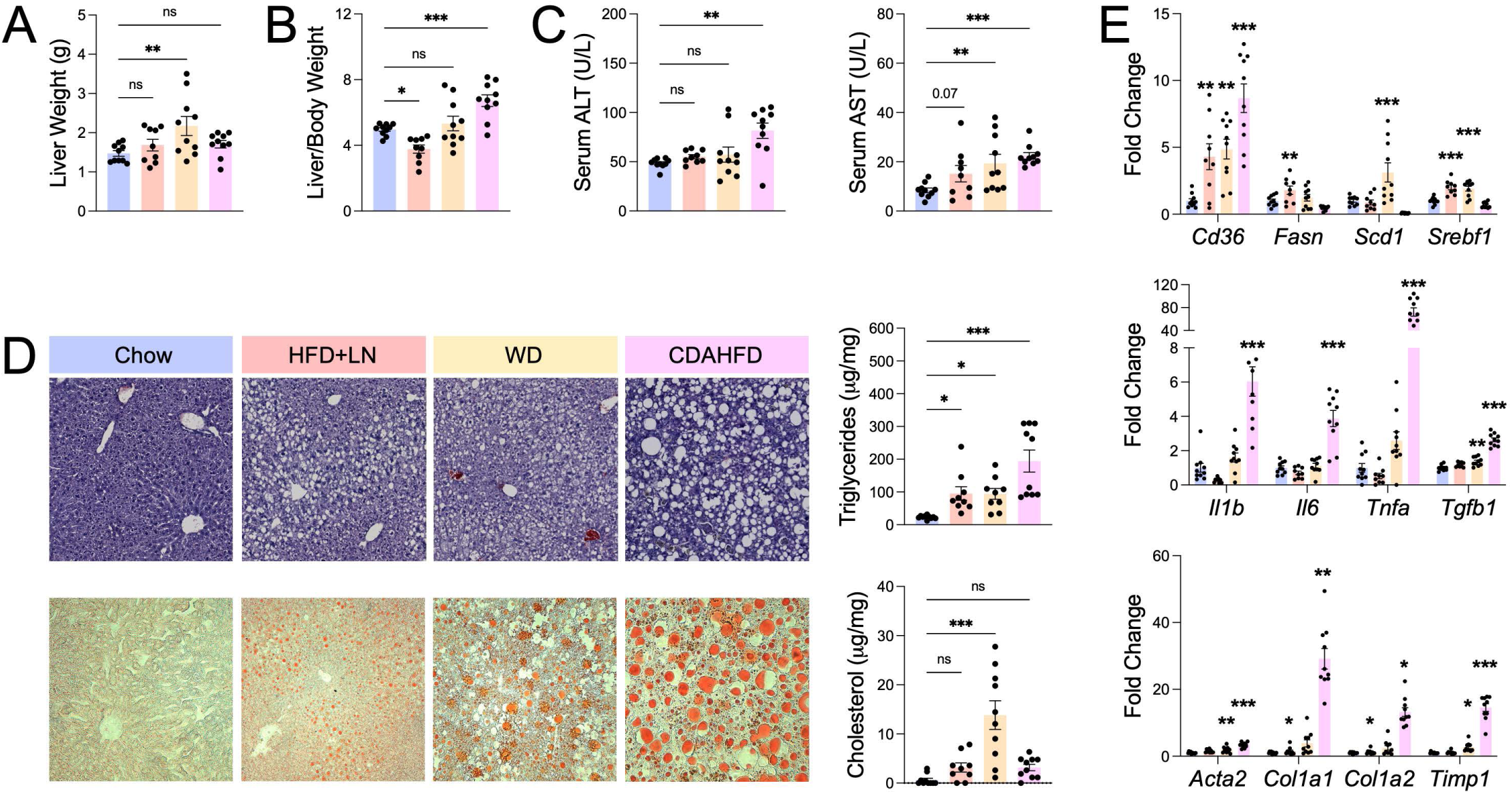
Hepatic steatosis, injury, and remodeling differ across complementary cardiometabolic stress models. Male C57BL/6 mice were treated for 10 weeks prior to tissue collection. **A** Liver weight. **B** Liver-to-body weight ratio. **C** Serum alanine aminotransferase (ALT) and aspartate aminotransferase (AST) concentrations. **D** Representative hematoxylin and eosin (H&E) and Oil Red O-stained liver sections with quantification of hepatic triglyceride and cholesterol content. **E** Hepatic gene expression of fatty acid metabolism, inflammatory, and fibrotic markers. Indicated *P* values represent comparison with chow controls. Data are presented as mean ± SEM. *n* = 9-10 mice/group pooled from two or more independent experiments. *\*P <* 0.05, **P < 0.01, ***P < 0.001 by one-way ANOVA followed by Tukey’s test.

### Cardiometabolic stress differentially influences cardiac structure and function

We therefore asked whether these distinct systemic and hepatic manifestations of cardiometabolic stress nevertheless converged on a shared cardiac phenotype. HFD+LN, WD, and CDAHFD each promoted myocardial fibrosis, as demonstrated by increased Picrosirius Red staining and induction of fibrotic genes, including *Acta2*, *Col1a1*, and *Timp1* (Figure 3A; Supplemental Figure 3A). Systolic function remained largely preserved, with no substantial reductions in ejection fraction or fractional shortening (Figure 3B). In contrast, Doppler analysis demonstrated impaired diastolic function, including changes in mitral E-wave and annular e′ velocities and increased E/e′, an index of elevated left ventricular filling pressure (Figure 3C). These changes were accompanied by prolonged isovolumetric relaxation time and mitral valve deceleration time. Global longitudinal strain was also impaired, particularly following HFD+LN treatment (Supplemental Figure 3B), consistent with subclinical systolic dysfunction and its reported association with myocardial stiffness in HFpEF (18).

**Figure 3.**
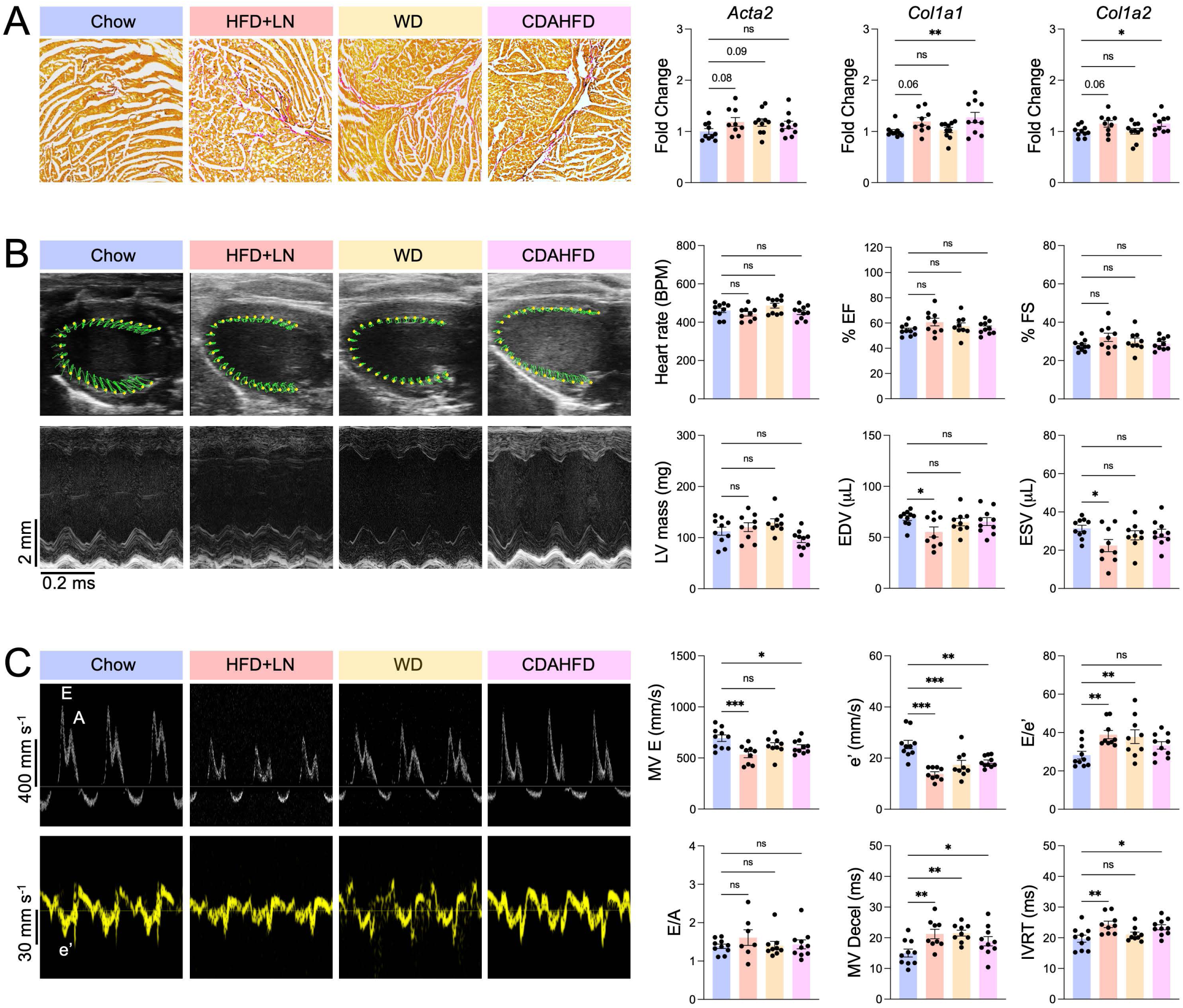
Complementary cardiometabolic stress models converge on cardiac fibrosis and diastolic dysfunction. Male C57BL/6 mice were treated for 10 weeks prior to assessments. **A** Representative Picrosirius Red-stained heart sections with quantification of cardiac fibrotic marker genes. **B** Echocardiographic assessment of systolic function, including heart rate beats per minute (BPM), ejection fraction (EF), fractional shortening (FS), left ventricular (LV) mass, end-diastolic volume (EDV), and LV end-systolic volume (ESV). **C** Echocardiographic assessment of diastolic function, including mitral valve deceleration time (MV decel), isovolumetric relaxation time (IVRT), E/A ratio, E/e’ ratio, and e’ velocity. Data are presented as mean ± SEM. *n* = 9-10 mice/group pooled from two or more independent experiments. *\*P <* 0.05, *\*\*P <* 0.01, ***P < 0.001 by one-way ANOVA followed by Tukey’s test.

Cardiac inflammatory responses were more model-dependent. HFD+LN increased expression of the macrophage-associated gene, including *Cd68* (Supplemental Figure 3A), consistent with macrophage expansion previously observed in experimental and human HFpEF (7,19). In contrast, WD and CDAHFD increased *Ccl2*, *Ccr2*, and *Cd68*, consistent with a broader monocyte/macrophage recruitment signature, although CDAHFD lacked broad induction of inflammatory genes such as *Il1b* and *Tnfa* despite its pronounced inflammatory phenotype in the liver (Supplemental Figure 3A). These divergent inflammatory responses accompanied myocardial fibrosis and dysfunction across all three models, identifying shared cardiac consequences of cardiometabolic stress despite distinct systemic and hepatic phenotypes.

### Liver transcriptomics identifies conserved and model-specific responses to metabolic stress

Given the divergent severity of liver injury across cardiometabolic models, we next performed bulk RNA sequencing to define corresponding hepatic transcriptional responses. CDAHFD produced the most pronounced hepatic phenotype and was therefore prioritized to characterize transcriptional remodeling associated with liver-predominant disease. Principal component analysis demonstrated marked separation of CDAHFD from control livers (Figure 4A), accompanied by substantially greater transcriptional remodeling compared with HFD+LN and WD (Figure 4B). Consistent with this dominant response, unsupervised clustering readily distinguished CDAHFD from control livers (Figure 4C). Differential expression analysis revealed broad transcriptional changes in CDAHFD livers (Figure 4D), with pathway enrichment identifying immune and inflammatory responses, including T cell activation and TNF production, together with cell death, response to wounding, and extracellular matrix organization (Figure 4E). At the gene level, CDAHFD suppressed hepatic metabolic genes, including *Cyp2c29*, *Ces2a*, *Aldh8a1*, and *Slc6a12*, while inducing genes associated with immune activation and tissue remodeling, including *Lgals3*, *Pla2g7*, *Sirpa*, *Lyz2*, and *Scara3* (Figure 4F). These findings demonstrate extensive inflammatory and tissue-remodeling programs accompanying the pronounced liver injury induced by CDAHFD.

**Figure 4.**
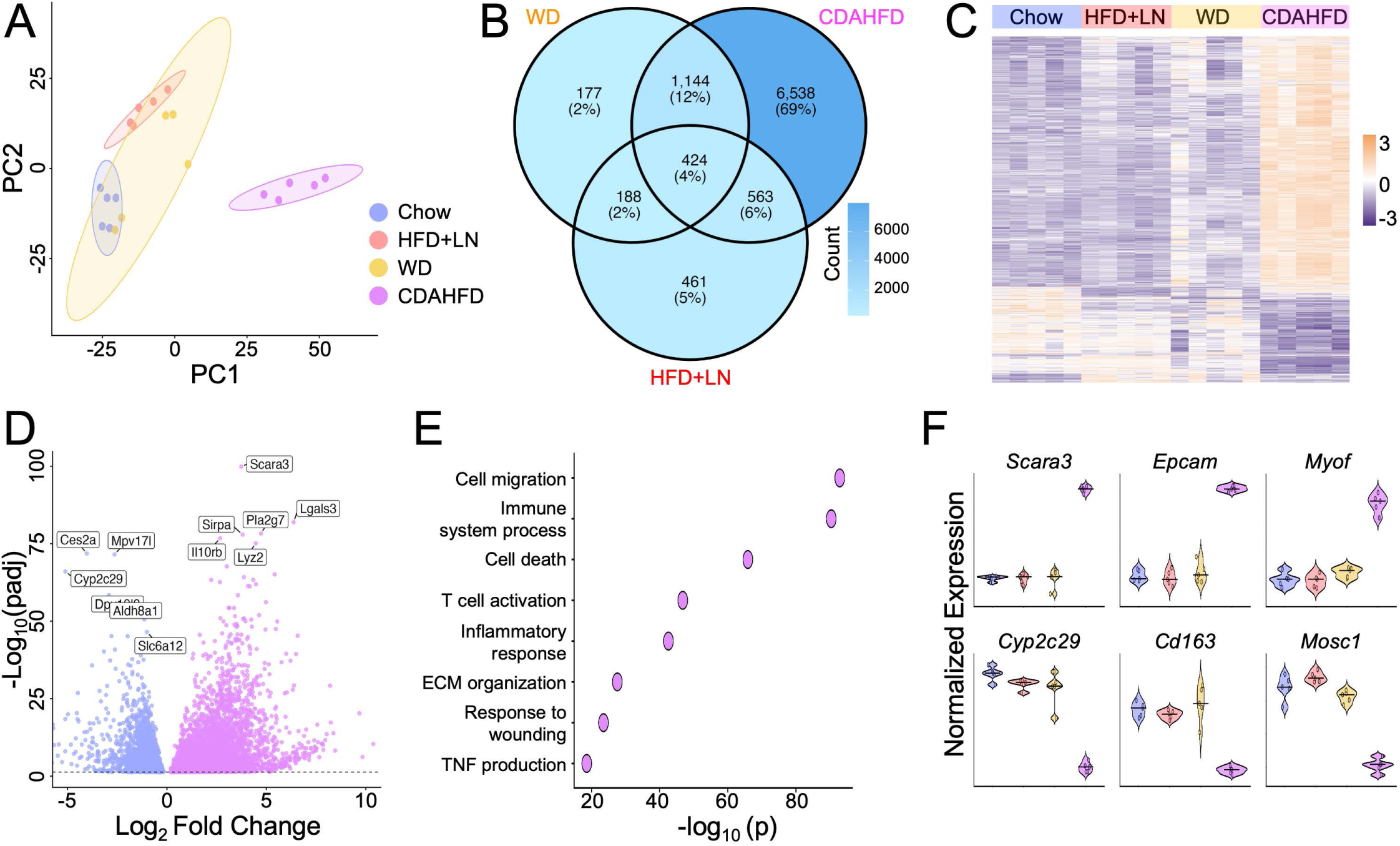
Hepatic transcriptomic profiling identifies model-specific responses to cardiometabolic stress. Male C57BL/6 mice were treated for 10 weeks prior to liver tissue collection for bulk RNA sequencing. **A** Principal component (PC) analysis of hepatic transcriptomes. **B** Venn diagram showing differentially expressed genes relative to chow controls. **C** Heatmap of differentially expressed genes. **D** Volcano plot of differential gene expression in CDAHFD versus chow. **E** Pathway enrichment analysis of differentially expressed genes upregulated in CDAHFD versus chow. **F** Expression of the three most upregulated and downregulated genes in CDAHFD compared with all experimental groups, *n* = 5 mice/group pooled from two independent experiments.

Although less extensive than CDAHFD, HFD+LN and WD produced distinct hepatic transcriptional responses. Unsupervised clustering showed comparatively modest global changes in both HFD+LN and WD relative to the marked separation of CDAHFD from chow (Figure 4C). HFD+LN nevertheless demonstrated clear separation from chow by principal component analysis (Figure 4A), indicating a distinct transcriptional state despite the more limited magnitude of differential gene expression. Pathway enrichment of HFD+LN livers centered on small molecule and fatty acid metabolism, lipid oxidation, ketone metabolism, and regulation of hormone levels, accompanied by increased expression of genes associated with metabolic and endocrine responses, including *Acadm* and *Serpina6* (Supplemental Figures 4A-C). In contrast, WD exhibited a more intermediate transcriptional profile characterized by immune and stress responses, including cell activation, phagocytosis, and response to lipid, together with changes in genes involved in lipid and cholesterol homeostasis, such as *Abcg5* and *Nrg4* (Supplemental Figures 4D-F). Thus, the intermediate WD response contrasted with the predominantly metabolic program induced by HFD+LN, while both remained less extensive than the inflammatory and remodeling response to CDAHFD. These findings define a spectrum of transcriptionally distinct hepatic states across cardiometabolic stress models.

We next asked whether common transcriptional responses emerged despite these model-specific differences in the magnitude and nature of hepatic injury. Pathway analysis of genes altered across all three models identified conserved regulation of responses to stimuli and cell communication together with small molecule metabolism, lipid binding, and catabolic processes (Supplemental Figure 5A). Examination of individual genes within this shared response revealed concordant induction of *Mfsd2a*, *Cyp4a14*, and *Themis* across HFD+LN, WD, and CDAHFD (Supplemental Figure 5B). Notably, each of these genes is similarly increased in human MASLD liver (20–22), supporting the translational relevance of this conserved hepatic response. These analyses identify a core molecular response maintained across otherwise distinct degrees and manifestations of hepatic cardiometabolic stress.

### Distinct cardiometabolic stressors induce model-specific cardiac transcriptional programs with conserved metabolic remodeling

Having established substantial model-dependent hepatic transcriptional remodeling, we asked whether the shared cardiac fibrosis and dysfunction observed across models were similarly accompanied by common or model-specific molecular responses. Bulk RNA sequencing demonstrated model-dependent separation of cardiac transcriptomes, while comparison of differentially expressed genes revealed both shared and model-specific responses across HFD+LN, WD, and CDAHFD (Figures 5A, 5B). As HFD+LN was developed as an experimental model of HFpEF (12), we first prioritized this condition to define HFpEF-associated cardiac transcriptional remodeling. HFD+LN produced broad transcriptional changes relative to chow (Figures 5C, 5D). Pathway enrichment identified lipid metabolism and fatty acid β-oxidation together with acyl-CoA metabolism, cardiac muscle contraction, and responses to endogenous and cellular stress (Figure 5E), consistent with metabolic and functional remodeling characteristic of human HFpEF (23,24). Among the most differentially expressed genes, *Erc1* was increased, whereas *Art3* was among the most strongly decreased genes in HFD+LN hearts (Figure 5F), consistent with its recently reported downregulation during pathological cardiac remodeling and heart failure (25).

**Figure 5.**
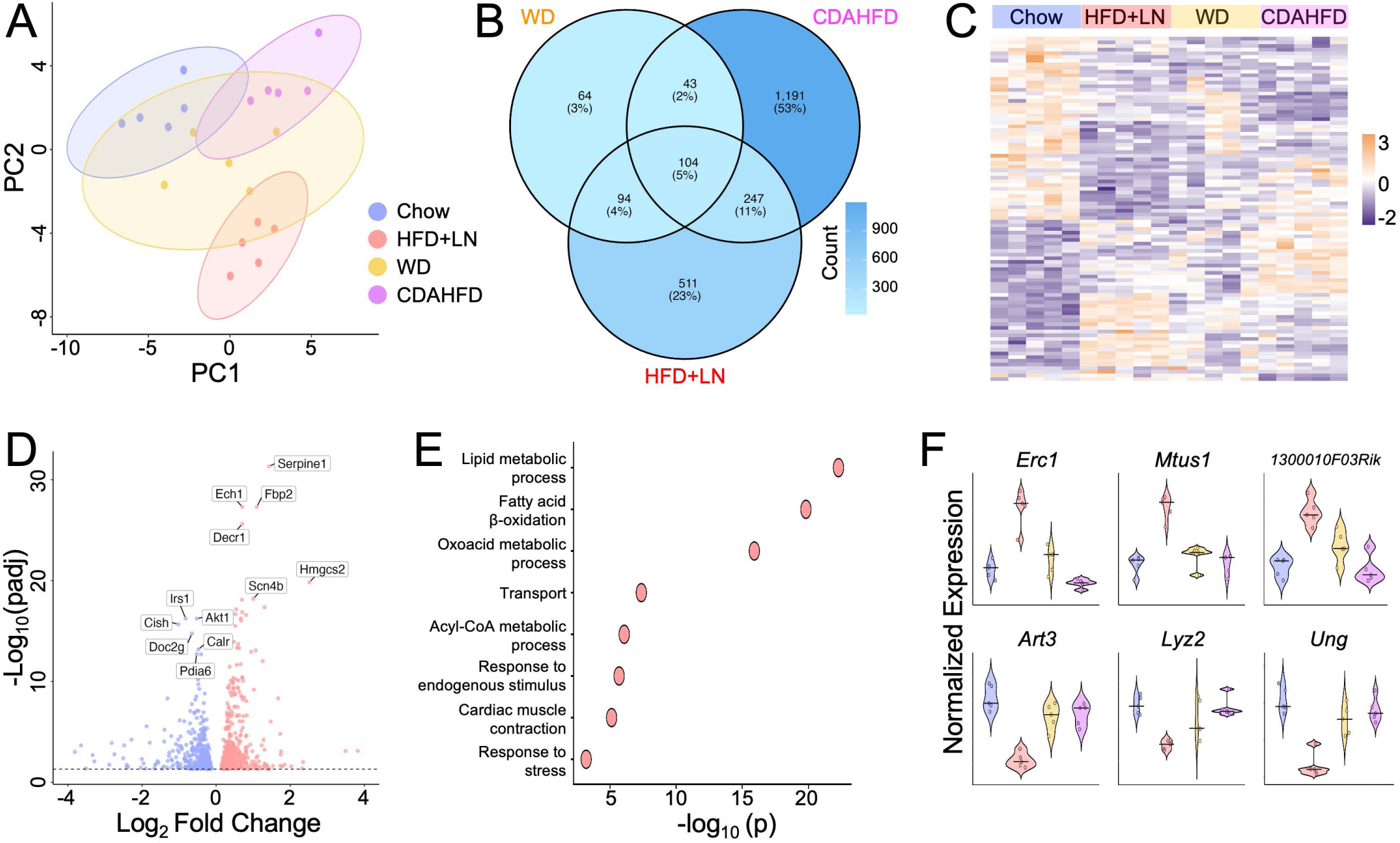
Cardiac transcriptomic profiling identifies model-specific responses to cardiometabolic stress. Male C57BL/6 mice were treated for 10 weeks prior to heart tissue collection for bulk RNA sequencing. **A** Principal component (PC) analysis of cardiac transcriptomes. **B** Venn diagram showing differentially expressed genes relative to chow controls. **C** Heatmap of differentially expressed genes. **D** Volcano plot of differential gene expression in HFD+LN versus chow. **E** Pathway enrichment analysis of differentially expressed genes upregulated in HFD+LN versus chow. **F** Expression of the three most upregulated and three most downregulated genes in HFD+LN compared with all experimental groups, *n* = 5 mice/group pooled from two independent experiments.

WD and CDAHFD similarly remodeled the cardiac transcriptome but produced distinct molecular signatures. WD exhibited a more intermediate transcriptional profile by unsupervised clustering (Figure 5C). Pathway analysis identified enrichment of fatty acid metabolism and β-oxidation, lipid transport and storage, and cholesterol metabolism, including changes in *Srebf1*, *Plin2*, and *Lpcat3* (Supplemental Figures 6A-C). These findings demonstrate substantial myocardial metabolic remodeling in the absence of the hypertension characteristic of HFD+LN or severe hepatic injury produced by CDAHFD. In contrast, CDAHFD demonstrated marked transcriptional separation from chow (Figure 5C) and unexpectedly produced the greatest number of cardiac differentially expressed genes, exceeding even HFD+LN (Figure 5B), despite its predominant hepatic phenotype. CDAHFD hearts were enriched for stress and defense responses, programmed cell death, cytokine responses, and leukocyte migration (Supplemental Figures 6D-F). Changes in *Gpr146*, *Pycard*, and *Pik3ip1* further implicated lipid homeostasis and inflammatory and immune regulation in this extensive cardiac response. These findings establish transcriptionally distinct cardiac responses across models despite their shared development of myocardial fibrosis and dysfunction.

Within these distinct transcriptional states, conserved changes centered predominantly on myocardial metabolism. Pathway analysis of genes altered across all three models identified shared enrichment of lipid metabolism, fatty acid β-oxidation, and lipid transport and localization, together with broader carboxylic acid and oxoacid metabolism (Supplemental Figure 7A). This conserved response was reflected at the gene level by concordant changes in *Acot1*, *Acsl1*, *Decr1*, and *Lpcat3*, which regulate complementary aspects of fatty acid activation, oxidation, and lipid remodeling (Supplemental Figure 7B). Together, these findings identify myocardial lipid metabolic remodeling as a core transcriptional response accompanying cardiac fibrosis and dysfunction across otherwise distinct manifestations of cardiometabolic stress.

### Integrated heart and liver transcriptomics identifies *Serpine1* as a conserved cardiometabolic stress response

To distinguish conserved molecular responses from organ-specific adaptations across cardiometabolic disease, we integrated heart and liver transcriptomes spanning HFD+LN, WD, and CDAHFD. Principal component analysis demonstrated dominant separation by tissue, with additional model-dependent variation within heart and liver samples (Figure 6A). Consistent with this distinction, hierarchical clustering identified organ- and disease-associated transcriptional programs enriched for inflammatory, metabolic, lipid biosynthetic, and extracellular matrix pathways (Figure 6B). The magnitude of transcriptional remodeling also varied substantially across the six model tissue comparisons, with the greatest number of differentially expressed genes observed in CDAHFD liver and considerably smaller responses in several cardiac conditions (Figure 6C). Given this heterogeneity, we reasoned that molecular responses maintained across these otherwise distinct conditions could identify core features of cardiometabolic disease. Despite thousands of differentially expressed genes across individual comparisons, UpSet analysis, which is designed for visualizing multi-set intersections (26), identified only five genes conserved across all six disease tissue conditions: *Serpine1*, *Ucp2*, *Acot2*, *Cep128*, and *Mylip* (Figure 6D). Examination of this highly restricted signature demonstrated consistent induction of *Serpine1* across models and tissues (Figure 6E), extending its independent identification among conserved hepatic and cardiac responses (Supplemental Figures 5B and 7B). Direct comparison confirmed increased *Serpine1* expression across HFD+LN, WD, and CDAHFD in both heart and liver (Figure 6F), identifying *Serpine1* as a conserved molecular response to cardiometabolic stress across disease context and target organ.

**Figure 6.**
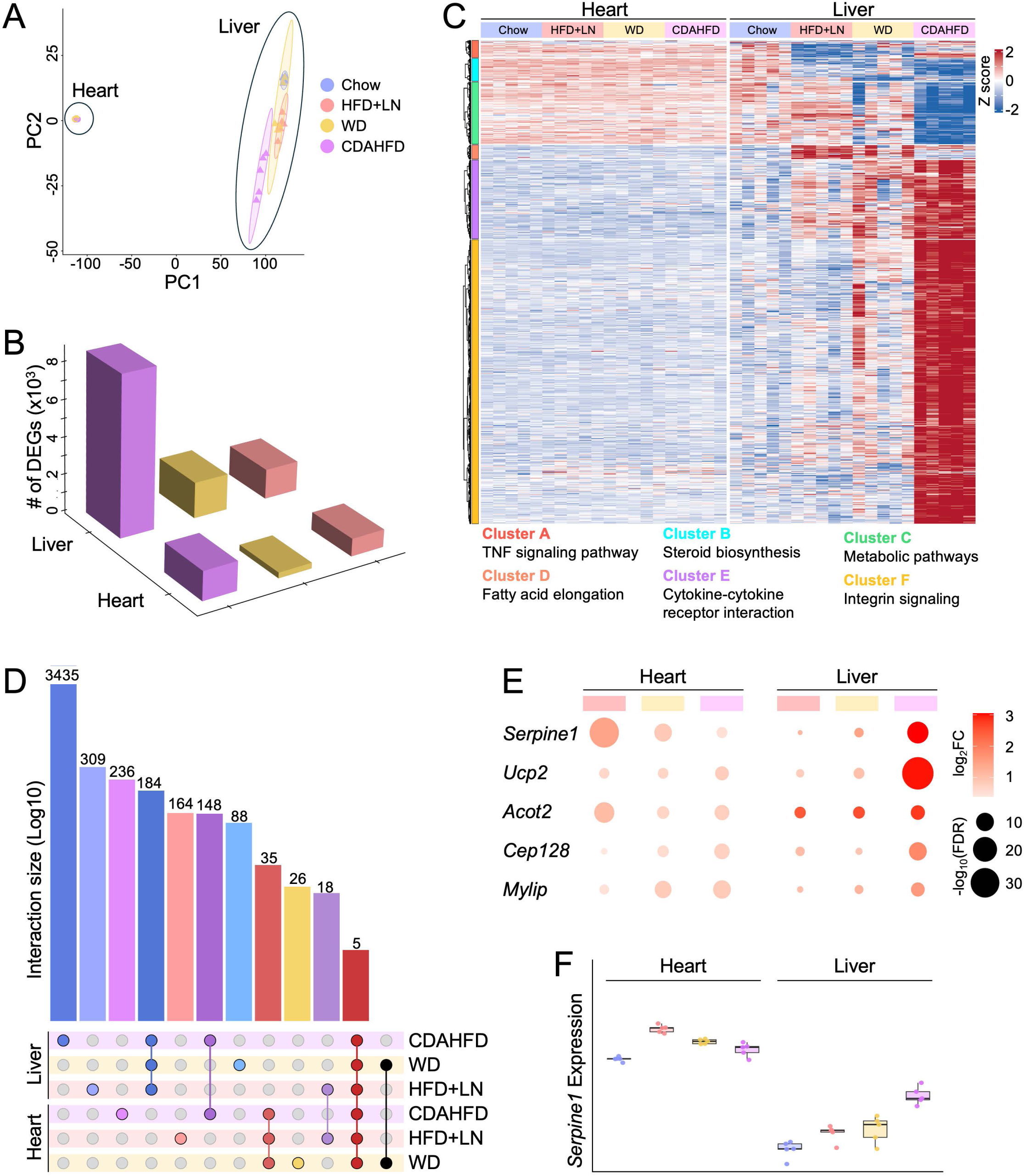
Integrated heart and liver transcriptomics identifies *Serpinel* as a conserved cardiometabolic stress response. Male C57BL/6 mice were treated for 10 weeks prior to heart and liver bulk RNA sequencing. HFD+LN, WD, and CDAHFD were compared with tissue-matched chow controls and integrated across models and organs. **A** Principal component (PC) analysis of heart and liver transcriptomes. **B** Heatmap of differentially expressed genes with representative transcriptional clusters and enriched pathways. **C** Three-dimensional representation of differentially expressed gene numbers across the six disease comparisons. **D** UpSet plot showing shared and condition-specific differentially expressed genes. **E** Bubble plot of the five most conserved upregulated genes. Color indicates log_2_ fold change and size indicates statistical significance (-logi_0_ false discovery rate). **F** *Serpinel* expression across heart and liver groups. Box-and-whisker plots show median, interquartile range, minimum and maximum values, and individual biological replicates. *n =* 5 mice/group pooled from two independent experiments.

**Figure 7.**
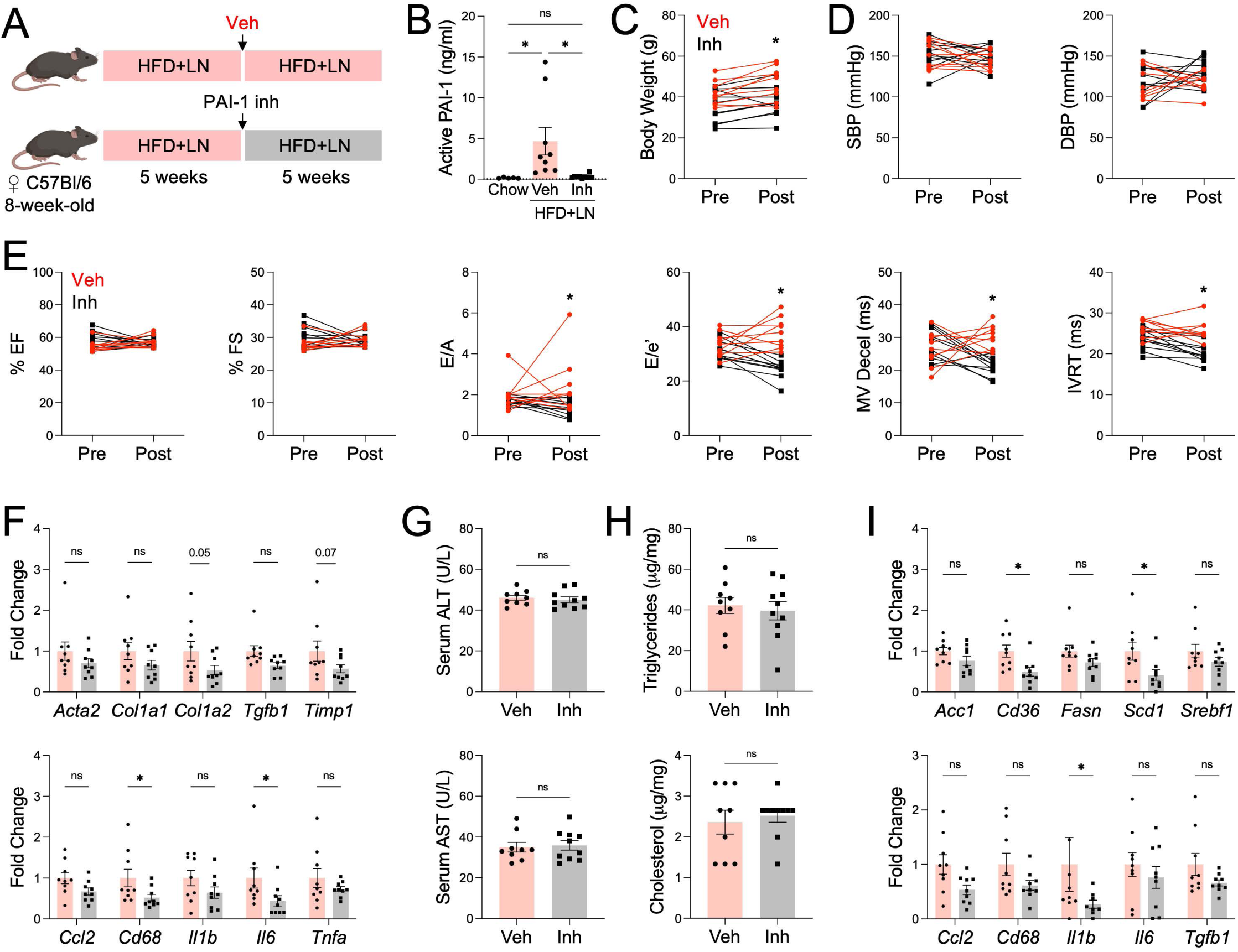
PAI-1 inhibition during established HFpEF improves cardiac remodeling and diastolic function. **A** Female C57BL/6 mice were treated with HFD+LN beginning at 8 weeks of age. Following establishment of HFpEF, mice were randomized to receive vehicle or PAI-1 inhibitor for an additional 5 weeks. **B** Serum active PAI-1 concentrations. **C** Body weight change. **D** Systolic (SBP) and diastolic (DBP) blood pressure. **E** Echocardiographic assessment of cardiac function, including ejection fraction (EF), fractional shortening (FS), mitral inflow E/A ratio, mitral E/e’ ratio, mitral valve deceleration time (MV decel), and isovolumetric relaxation time (IVRT). **F** Cardiac expression of inflammatory and fibrotic marker genes. **G** Serum alanine aminotransferase (ALT) and aspartate aminotransferase (AST) concentrations. **H** Hepatic triglyceride and cholesterol content. **I** Hepatic expression of fatty acid metabolism and inflammatory genes. Data are presented as mean ± SEM. *n* = 9-10 mice per group pooled from two independent experiments. *\*P <* 0.05 by unpaired *t* test one-way ANOVA followed by Tukey’s test.

### PAI-1 inhibition reverses cardiac dysfunction in established HFpEF without substantially altering hepatic injury

*Serpine1* encodes plasminogen activator inhibitor-1 (PAI-1), a circulating protein associated with metabolic dysfunction and fibrosis and, notably, with incident HFpEF and adverse outcomes in patients with HFpEF (27–29). Therefore, we tested whether pharmacologic PAI-1 inhibition could modify disease after HFpEF was established. Female C57BL/6 mice were treated with HFD+LN for 5 weeks to establish weight gain, hypertension, and diastolic dysfunction and then randomized to vehicle or PAI-1 inhibitor while HFD+LN treatment continued for an additional 5 weeks (Figure 7A). PAI-1 inhibition reduced circulating active PAI-1 (Figure 7B) and blunted further weight gain without substantially altering systolic or diastolic blood pressure (Figures 7C, 7D). Systolic function remained preserved, whereas PAI-1 inhibition improved multiple indices of diastolic function, including E/e′, mitral valve deceleration, and isovolumetric relaxation time (Figure 7E). These functional improvements were accompanied by reduced cardiac inflammatory and fibrotic gene expression (Figure 7F), consistent with attenuation of pathological myocardial remodeling. In the liver, PAI-1 inhibition did not substantially alter serum ALT or AST levels or hepatic lipid accumulation (Figures 7G, 7H). However, PAI-1 inhibition attenuated HFpEF-associated increases in the fatty acid metabolism genes *Cd36* and *Scd1* and the inflammatory gene *Il1b*, while *Ccr2* and *Cd68* were also reduced, albeit non-significantly (Figure 7I). Collectively, our comparative cross-model and cross-organ approach identified conserved molecular responses across otherwise distinct manifestations of cardiometabolic stress, ultimately revealing PAI-1 as a therapeutically actionable mediator of cardiac remodeling and diastolic dysfunction in HFpEF.

## DISCUSSION

In this study, we leveraged three complementary models representing cardiovascular, systemic metabolic, and hepatic manifestations of cardiometabolic stress to distinguish conserved molecular responses from organ- and disease-specific adaptations. Despite distinct systemic and hepatic phenotypes, all three models developed cardiac fibrosis and dysfunction, while transcriptomic profiling revealed substantial heterogeneity across models and tissues. Integrated analysis distilled these diverse responses to only five genes conserved across all six heart and liver disease conditions, among which *Serpine1* emerged as a consistently upregulated response to cardiometabolic stress. Importantly, pharmacologic inhibition of its encoded protein, PAI-1, during established HFpEF improved cardiac remodeling and diastolic function, with more limited effects on hepatic injury and steatosis. Collectively, these findings demonstrate the utility of comparative cross-model and cross-organ analyses to uncover conserved disease biology and identify PAI-1 as a therapeutically actionable mediator of cardiac dysfunction during HFpEF.

Cardiometabolic diseases rarely occur in isolation, yet the molecular responses shared across their diverse manifestations remain poorly defined. Our models captured this heterogeneity, with distinct systemic metabolic profiles and a gradient of hepatic injury ranging from predominantly metabolic adaptation to inflammation and fibrosis. Notably, circulating metabolites altered across our experimental models overlapped with metabolic abnormalities reported in human HFpEF and MASLD (30), supporting the translational relevance of these distinct metabolic phenotypes. Inclusion of WD was particularly informative because it provided an intermediate model of systemic metabolic stress without the hypertension characteristic of HFD+LN or severe hepatic remodeling induced by CDAHFD. The hepatic transcriptional response further emphasized that conserved changes need not uniformly represent pathogenic signaling. *Mfsd2a*, *Themis*, and *Cyp4a14* were increased across models and in human MASLD (20–22). However, functional studies suggest divergent roles with MFSD2A-dependent lysophospholipid uptake and THEMIS protecting against steatotic liver disease, whereas CYP4A14 promotes steatosis, inflammation, and fibrosis. Similarly, high fat diet-associated induction of *Nrg4* may represent an adaptive response, consistent with recent evidence that NRG4 restrains hepatic inflammatory signaling in MASLD (31). These observations underscore an important feature of comparative analyses where conserved and model-specific responses can encompass both compensatory and pathogenic pathways, requiring functional interrogation to distinguish adaptation from disease drivers.

In contrast to the heterogeneous hepatic response, myocardial fibrosis and diastolic dysfunction emerged across all three models, suggesting that distinct cardiometabolic insults can arrive at a similar cardiac phenotype. This finding is consistent with the recognized relationship between MASLD and cardiovascular dysfunction, including evidence that steatohepatitis itself can promote diastolic dysfunction in aged mice (32). Impaired global longitudinal strain, particularly in HFD+LN, provided additional evidence of myocardial dysfunction and parallels its prognostic significance in human HFpEF and association with subclinical cardiac dysfunction in MASLD (33,34). Despite this phenotypic overlap, cardiac transcriptomes remained highly model dependent. HFD+LN prominently altered fatty acid metabolic pathways, consistent with metabolic remodeling reported in human HFpEF, while individual gene changes further connected this response to broader mechanisms of cardiac disease. *Art3* was decreased in HFD+LN hearts, consistent with its reported downregulation in pressure overload-induced heart failure (25), whereas increased *Erc1* was notable given human genetic associations between the *ERC1* locus and coronary microvascular function (35), a pathway strongly implicated in HFpEF (36). WD and CDAHFD engaged distinct transcriptional programs, with CDAHFD unexpectedly producing the most extensive cardiac response despite its liver-predominant phenotype. Against this model-specific heterogeneity, conserved lipid metabolic responses across all three conditions identify myocardial metabolic remodeling as a shared feature of cardiometabolic cardiac stress, although the dietary lipid exposure common to these models likely contributes to the prominence of these pathways.

Integration across heart and liver provided an opportunity to move beyond these model- and organ-specific responses. Recent systems-biology studies have begun to identify candidate mediators of liver-heart communication in HFpEF, highlighting the potential importance of interorgan signaling in cardiometabolic disease (11). Our findings similarly revealed potentially coordinated responses across tissues. For example, the CDAHFD liver exhibited prominent TNF-associated inflammatory signaling and the CDAHFD heart was enriched for cytokine-response pathways, raising the possibility of interorgan crosstalk in which the liver serves as a source of inflammatory signals received by the heart. However, establishing such relationships remains challenging because coordinated molecular changes cannot define the direction or source of interorgan communication. Similarly, an intervention that improves hepatic metabolism and cardiac function does not necessarily establish liver-heart signaling, as the targeted pathway may act locally in both organs or improved hepatic health may indirectly reduce systemic stress on the heart. Rather than attempting to infer these relationships, our comparative strategy leveraged this complexity to ask which molecular responses persisted across multiple disease contexts and both organs, reducing thousands of transcriptional changes to only five conserved genes. This stringent intersection increases confidence that these signals reflect fundamental responses to cardiometabolic stress rather than consequences of a single model or tissue. *Serpine1*, which encodes PAI-1, was particularly compelling given its established links to obesity, insulin resistance, visceral adiposity, cellular senescence, and biological aging (28,37). PAI-1 is also one of the few circulating biomarkers uniquely associated with incident HFpEF but not HFrEF (38), while elevated tissue plasminogen activator/PAI-1 complexes independently predict cardiovascular and all-cause mortality in HFpEF (29). Human *SERPINE1* loss-of-function is associated with lower fasting insulin, reduced diabetes prevalence, and greater longevity (39), further connecting PAI-1 to the metabolic and aging biology characteristic of HFpEF.

Our therapeutic studies extended these associations by demonstrating that PAI-1 remains actionable after HFpEF is established. Previous work showed that the PAI-1 inhibitor, TM5441, attenuates LN-induced hypertension, cardiac hypertrophy, vascular fibrosis, and senescence when inhibition is initiated concurrently with LN exposure (27). In contrast, we initiated PAI-1 inhibition only after development of the HFpEF phenotype and observed improved diastolic function and reduced cardiac inflammatory and fibrotic remodeling without normalization of blood pressure. This distinction supports a therapeutic effect on established cardiac disease rather than attenuation of the initiating cardiometabolic stress. The comparatively modest hepatic response is also consistent with recent studies showing that PAI-1 deletion produces limited effects on hepatic lipid accumulation during high-fat diet feeding (40), suggesting that the consequences of PAI-1 signaling may differ across target organs. Together, these findings extend the role of PAI-1 from a biomarker and mediator of cardiometabolic aging to a therapeutically actionable pathway in established HFpEF and demonstrate how cross-model, cross-organ discovery can prioritize conserved signals with functional relevance.

Several limitations should be acknowledged. Although the comparative studies were predominantly performed in male mice, the therapeutic efficacy of PAI-1 inhibition was independently validated in female mice, supporting efficacy across sexes. Future studies directly comparing sex-specific molecular responses remain warranted given the higher prevalence of HFpEF among women (41). Studies were also performed in young adult mice, whereas the incidence of HFpEF and MASLD increases substantially with aging (42), warranting future studies to determine whether the conserved molecular responses and therapeutic effects of PAI-1 inhibition are maintained with aging. Finally, while the models used capture distinct manifestations of cardiometabolic disease, bulk RNA sequencing cannot resolve the cellular origins of these responses or distinguish tissue-specific effects from interorgan crosstalk. Future tissue- and cell-specific studies will be needed to resolve these mechanisms.

In summary, cardiometabolic disease is inherently heterogeneous, with overlapping systemic, hepatic, and cardiovascular manifestations that are difficult to capture within any single experimental model. Integrating complementary models across target organs provides a systems-level strategy to distinguish context-specific adaptations from conserved responses with greater potential to generalize across human disease. Our identification and therapeutic validation of PAI-1 in established HFpEF demonstrates how this approach can prioritize conserved biology and accelerate cardiovascular therapeutic discovery.

## AUTHOR CONTRIBUTIONS

M.S., H.J.J., B.R.L., S.P., and R.S. performed experiments, acquired data, and analyzed results. C.L., M.J.H., A.S., and Y.A. contributed to data analysis and interpretation. K.H.K. and M.D. conceived and designed the study, supervised the research, interpreted the data, and wrote the manuscript. All authors contributed to interpretation of the data, critically revised the manuscript, and approved the final version. M.S., H.J.J., and B.R.L. contributed equally to this work. K.H.K. and M.D. jointly supervised this work.

## ACKNOWLEDGMENTS

This work was supported by the National Institutes of Health (grant R01DK126656 to KHK), American Heart Association (25TPA1463689 to MD), and Institute for Perioperative Medicine (to KHK and MD). These studies used the Small Animal Cardiovascular Phenotyping Service Center at UTHealth-Houston and metabolomics services provided by the Metabolomics Facility at MD Anderson Cancer Center (supported in part by the University of Texas MD Anderson Cancer Center and P30CA016672).

## DATA AVAILABILITY

The raw bulk RNA-sequencing data generated in this study have been deposited in the NCBI Gene Expression Omnibus (GEO) under accession number GSE345186.

## METHODS

### Mice

Male and female C57BL/6J mice were purchased from Jackson Laboratory (Stock no. 000664) and housed in a temperature-and humidity-controlled specific pathogen-free facility on a 12-hour light/dark cycle with ad libitum access to food and water. Mice were 8 weeks of age at the initiation of experimental diets. All animal studies were approved by the Institutional Animal Care and Use Committee at University of Texas Health Science Center at Houston.

### Cardiometabolic Disease Models

Three complementary mouse models representing distinct manifestations of cardiometabolic stress were studied: high-fat diet plus L-NAME (HFD+LN)-induced heart failure with preserved ejection fraction (HFpEF), Western diet (WD)-induced obesity, and choline-deficient, L-amino acid-defined, high-fat diet (CDAHFD)-induced steatotic liver disease. HFD+LN was induced by feeding mice a 60% kcal high-fat diet (Research Diets, D12492) together with drinking water supplemented with 0.5 g/L L-NAME (Sigma-Aldrich, N5751) for 10 weeks as previously described (7). Diets and drinking water were replaced twice weekly, and age-matched chow-fed mice served as controls. Body weight was monitored weekly throughout the study. For the WD model, mice were fed a WD containing 40 kcal% fat, 35% sucrose, and 1.25% cholesterol (Research Diets, D12079B) for 10 weeks, whereas control animals remained on standard laboratory chow. Steatotic liver disease was induced by feeding mice a choline-deficient, L-amino acid-defined, high-fat diet (Research Diets, A06071302) for 10 weeks according to established protocols (43). This model produces hepatic steatosis, inflammation, and fibrosis with minimal obesity.

### Noninvasive Blood Pressure

Blood pressure was measured in conscious mice using the tail-cuff method with a CODA system (Kent Scientific). To minimize stress-related variability, mice were acclimated to the restrainer and tail-cuff apparatus on a heated platform (34-36°C) for 10-15 minutes per day over 3 consecutive days prior to data collection. On the day of measurement, mice were placed in acrylic restrainers on the warmed platform to promote tail vasodilation. Each recording session consisted of 10 acclimation cycles followed by 30 measurement cycles, with movement artifacts automatically excluded by the software. Systolic, diastolic, and mean arterial pressures, together with heart rate, were calculated as the average of at least 10 valid measurements. All recordings were performed at the same time of day to minimize circadian variation, and data represent the mean of 2-3 independent sessions per animal.

### PAI-1 Inhibitor Treatment

To determine the therapeutic contribution of PAI-1 following disease establishment, HFpEF mice were maintained on HFD+LN for 5 weeks, at which point the HFpEF phenotype (weight gain, hypertension, and diastolic dysfunction) was confirmed. Mice were then randomized to receive vehicle (corn oil) or the selective PAI-1 inhibitor TM5441 (20 mg/kg; MedChemExpress) administered by oral gavage twice weekly while continuing HFpEF treatment for an additional 5 weeks. Investigators performing molecular analyses were blinded to treatment assignment.

### Echocardiography

Cardiac structure and function were evaluated by transthoracic echocardiography at the indicated time points using a Vevo 3100 high-frequency ultrasound imaging system (FUJIFILM VisualSonics). Mice were anesthetized with 1-2% isoflurane delivered in 100% oxygen, and anesthetic depth was adjusted to maintain heart rates between 400 and 500 beats/min to preserve physiological cardiac function. Parasternal short-axis M-mode images were acquired at the mid-papillary level to measure left ventricular internal dimensions at end-diastole and end-systole from three consecutive cardiac cycles, from which left ventricular ejection fraction and fractional shortening were calculated using Vevo LAB software (FUJIFILM VisualSonics). Left ventricular global longitudinal strain was quantified from parasternal long-axis B-mode cine loops using VevoStrain software (FUJIFILM VisualSonics), with values averaged across three consecutive cardiac cycles. Diastolic function was assessed from the apical four-chamber view using pulse-wave Doppler to measure transmitral early (E) and late (A) filling velocities together with tissue Doppler imaging of the mitral annulus to quantify early diastolic myocardial velocity (e’) and additional indices of mitral valve function.

### Serum Biochemical Analyses

At the study endpoint (10 weeks), whole blood was collected by cardiac puncture immediately prior to euthanasia, and serum was isolated by centrifugation. Serum alanine aminotransferase (TECO Diagnostics), aspartate aminotransferase (TECO Diagnostics), triglycerides (Thermo Fisher), and total cholesterol (Thermo Fisher) were measured using commercially available colorimetric assay kits according to the manufacturers’ instructions.

### Serum PAI-1 Measurements

Circulating active PAI-1 concentrations were measured in mouse serum using the Mouse Active PAI-1 ELISA Kit (Innovative Research) according to the manufacturer’s instructions.

### Serum Metabolomics

Untargeted metabolomic profiling and targeted fatty acid analyses were performed on mouse serum using separate mass spectrometry-based platforms. For untargeted metabolomics, approximately 50-70 μL of serum per mouse was extracted with ice-cold 80:20 (v/v) methanol/water containing 0.1% ammonium hydroxide, as previously described (44). Extracts were centrifuged at 17,000 x *g* for 5 minutes at 4°C, and supernatants were evaporated to dryness under nitrogen and reconstituted in deionized water. Samples were analyzed by ion chromatography-high-resolution mass spectrometry (IC-HRMS) using a Dionex ICS-6000+ system equipped with an IonPac AS11 column coupled to a Thermo Orbitrap IQ-X Tribrid mass spectrometer operating in negative electrospray ionization mode. Metabolite abundances were normalized prior to principal component and differential metabolite analyses to define circulating metabolic signatures across experimental groups. Targeted profiling of circulating fatty acids was performed using a separate mass spectrometry-based platform, with relative abundances of individual fatty acid species compared across experimental groups.

### Histology

Liver and heart tissues were harvested at study completion and drop fixed in 4% PFA overnight at 4°C. Following fixation, tissues were rinsed in PBS for 2 hours and subsequently cryoprotected in PBS with 30% sucrose at 4°C for 48-72 hours. Liver and heart tissues were embedded in Tissue-Tek O.C.T. compound (Sakura) and cryosectioned at 10 μm thickness using a Leica CM1860 UV cryostat. Liver morphology was evaluated by hematoxylin and eosin (H&E) staining. Hepatic lipid accumulation was assessed by Oil Red O staining, with sections incubated in Oil Red O solution for 15 minutes, and myocardial fibrosis was assessed by Picrosirius Red staining, with sections incubated in Picrosirius Red solution for 1 hour. Stained tissue sections were imaged using an Aperio slide scanner (Leica). Histological analyses were performed by investigators blinded to treatment group.

### Bulk RNA Sequencing

Total RNA was isolated from heart and liver tissue using standard methods. All RNA integrity test, library preparation, and sequencing procedures were performed by Novogene Corporation (multi-omics facility in Beaverton, Oregon). RNA integrity was assessed using an RNA integrity number (RIN), and only samples with RIN values greater than 7 were used for sequencing. Total RNA was processed for mRNA library preparation using poly-A enrichment, and sequencing was executed on the Illumina NovaSeq X Plus platform, which generated 150 bp pair-end (PE150) read, yielding approximately 6 Gb of raw data per sample. Raw data underwent quality control filtering to remove low-quality reads and adapter sequences prior to downstream bioinformatics analysis.

### Transcriptomic Data Processing and Bioinformatic Analyses

FASTQ files containing raw sequencing reads were obtained from Novogene, and sequence quality was assessed using FastQC (v0.12.1). Reads were then aligned to the Mus musculus reference genome (GRCm38/mm10) using HISAT2, followed by alignment processing with SAMtools. Gene expression was subsequently quantified using featureCounts to generate a raw count matrix across all samples. The resulting counts matrix was imported into R for downstream analyses. Genes with low expression were filtered prior to downstream analysis, retaining genes with at least 10 counts in at least three samples. Normalized counts were obtained using the DESeq2 variance-stabilizing transformation (vst) to correct for sequencing depth and biases. Principal component analysis (PCA) was performed on vst-normalized counts to assess sample clustering. Differential expression analysis was carried out in DESeq2 using the Wald test with Benjamini–Hochberg correction. Genes were considered differentially expressed if they exhibited an adjusted p-value less than 0.05. Volcano plots were generated to visualize top differentially expressed genes (DEGs) in each condition, and heatmaps to visualize relative expression patterns of DEGs across all samples. Pathway enrichment analysis was performed by g:Profiler (v0.2.4) on significant DEGs identified between each condition and the control groups. Genes were separated by direction of change of the log2 fold-change and an adjusted p-value less than 0.05 was used to identify significant pathways enriched in each group. Lollipop plots were generated to visualize the significance of top enriched pathways. Visualizations were generated using ggplot2 (v4.0.3) and pheatmap (v1.0.13).

### Integrated Transcriptomic Analyses

Heart and liver transcriptomic datasets were analyzed independently and subsequently integrated to identify organ-specific and conserved molecular responses across complementary manifestations of cardiometabolic stress. For the integrated analysis, count data were transformed using the DESeq2 variance-stabilizing transformation (VST), and principal component analysis (PCA) was performed on VST-normalized counts from a DESeq2 dataset designed to account for both organ and condition to assess sample clustering. Differential expression analysis was then performed separately for each organ using DESeq2 with the Wald test and Benjamini–Hochberg correction for multiple comparisons. Genes with an adjusted p-value < 0.05 were considered differentially expressed. Significant genes from the heart and liver analyses were then combined into a shared dataframe, with genes included if they reached the significance threshold in at least one of the six experimental comparisons. An UpSet plot was generated from the shared dataset to visualize overlap among upregulated differentially expressed genes across the six comparisons and identify conserved gene expression responses across groups. Differentially expressed genes shared across all groups were identified and further investigated. A combined heatmap was subsequently generated to visualize relative expression patterns of DEGs across all heart and liver samples. For heatmap visualization, genes were filtered using more stringent criteria of an adjusted p-value < 0.01 and an absolute log2 fold change > 1. Hierarchical clustering was performed using Euclidean distance and Ward.D2 linkage, and the resulting dendrogram was divided into six clusters. Pathway enrichment analysis was subsequently performed separately for each of the six clusters to identify biological pathways associated with each distinct expression profile.

### Quantitative Polymerase Chain Reaction

Total RNA was isolated from heart and liver tissue using TRIzol reagent and reverse transcribed into cDNA using iScript Reverse Transcription Supermix (BIO-RAD). Quantitative PCR was performed using SYBR Green chemistry on a BIO-RAD CFX Opus 384 Real-Time PCR System. Relative gene expression was calculated using the ΔΔCt method following normalization to an endogenous housekeeping gene. Primer sequences are provided in the resources table.

### Statistical Analysis

Statistical analyses were performed using GraphPad Prism 10 (GraphPad Software). Data are presented as mean ± SEM. Comparisons between two groups were performed using an unpaired two-tailed Student’s *t* test. Comparisons involving more than two groups were analyzed by one-way or two-way ANOVA followed by Tukey’s multiple-comparison post hoc test, as appropriate. Differential gene expression analyses incorporated Benjamini-Hochberg correction for multiple testing. Statistical significance was defined as a two-sided P value < 0.05.

## Notes

### Competing Interest Statement

The authors have declared no competing interest.

