## Supplemental Figures for "Complementary Models of Cardiometabolic Stress Reveal Conserved Molecular Programs Driving Cardiac Remodeling"

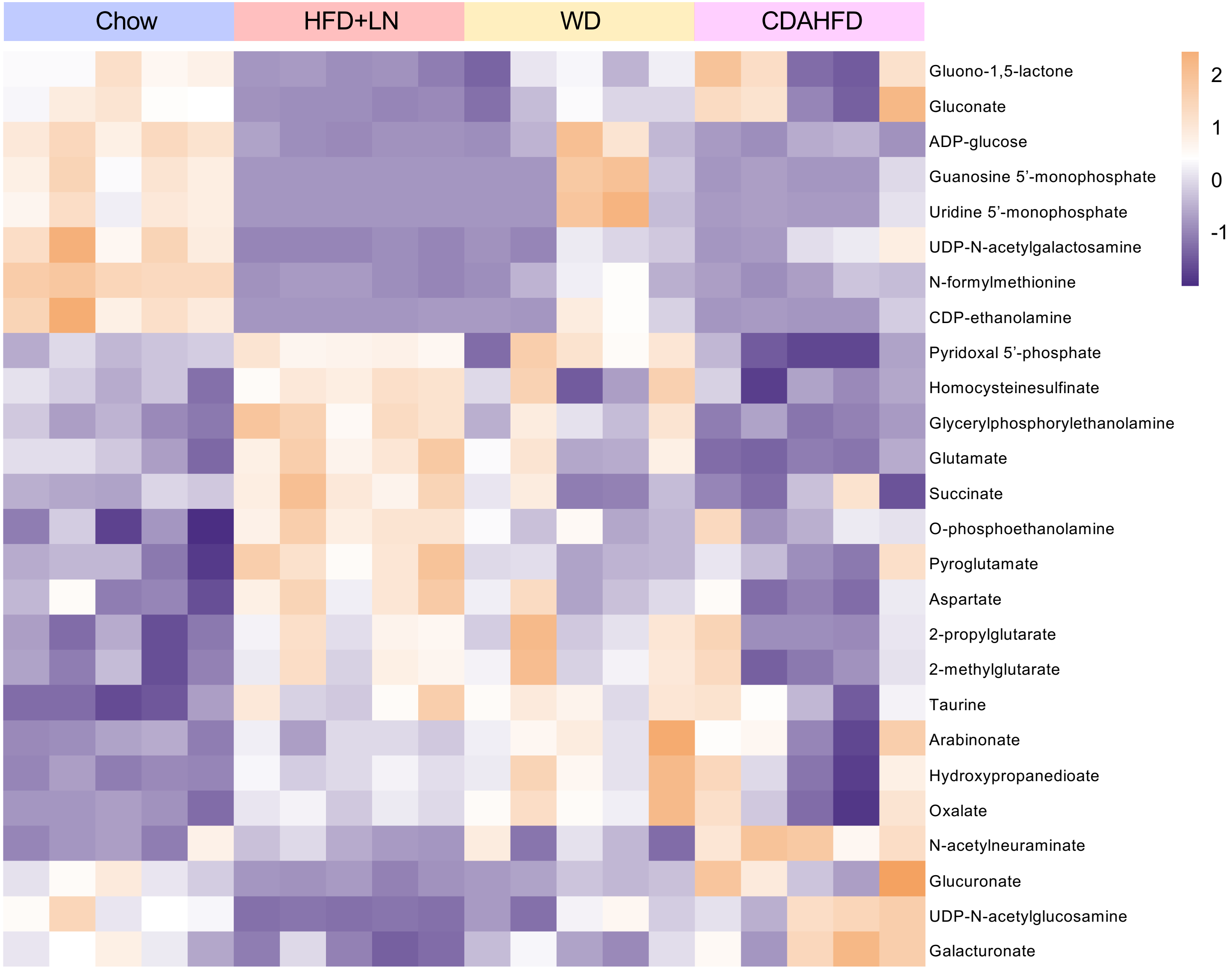


**Supplemental Figure 1. Complementary cardiometabolic stress models produce distinct circulating metabolomic signatures.** Male C57BL/6 mice were treated for 10 weeks prior to serum collection. Heatmap showing differentially abundant serum metabolites across chow, HFD+LN, WD, and CDAHFD groups. Metabolite abundances are displayed as row-scaled normalized values. *n* = 5 mice/group pooled from two independent experiments.

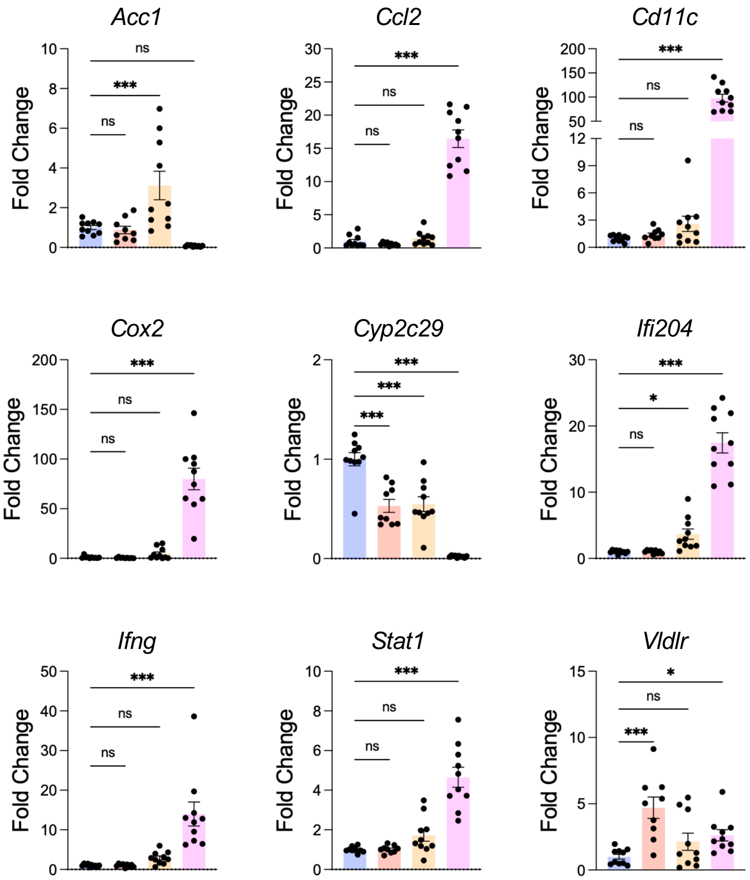


**Supplemental Figure 2. Hepatic gene expression differs across complementary cardiometabolic stress models.** Male C57BL/6 mice were treated for 10 weeks prior to liver tissue collection. Hepatic gene expression of markers associated with fatty acid metabolism, inflammation, and fibrosis. Data are presented as mean ± SEM. $n$ = 9–10 mice per group pooled from two or more independent experiments. *$P < 0.05$, ***$P < 0.001$ by one-way ANOVA followed by Tukey's multiple comparisons test.

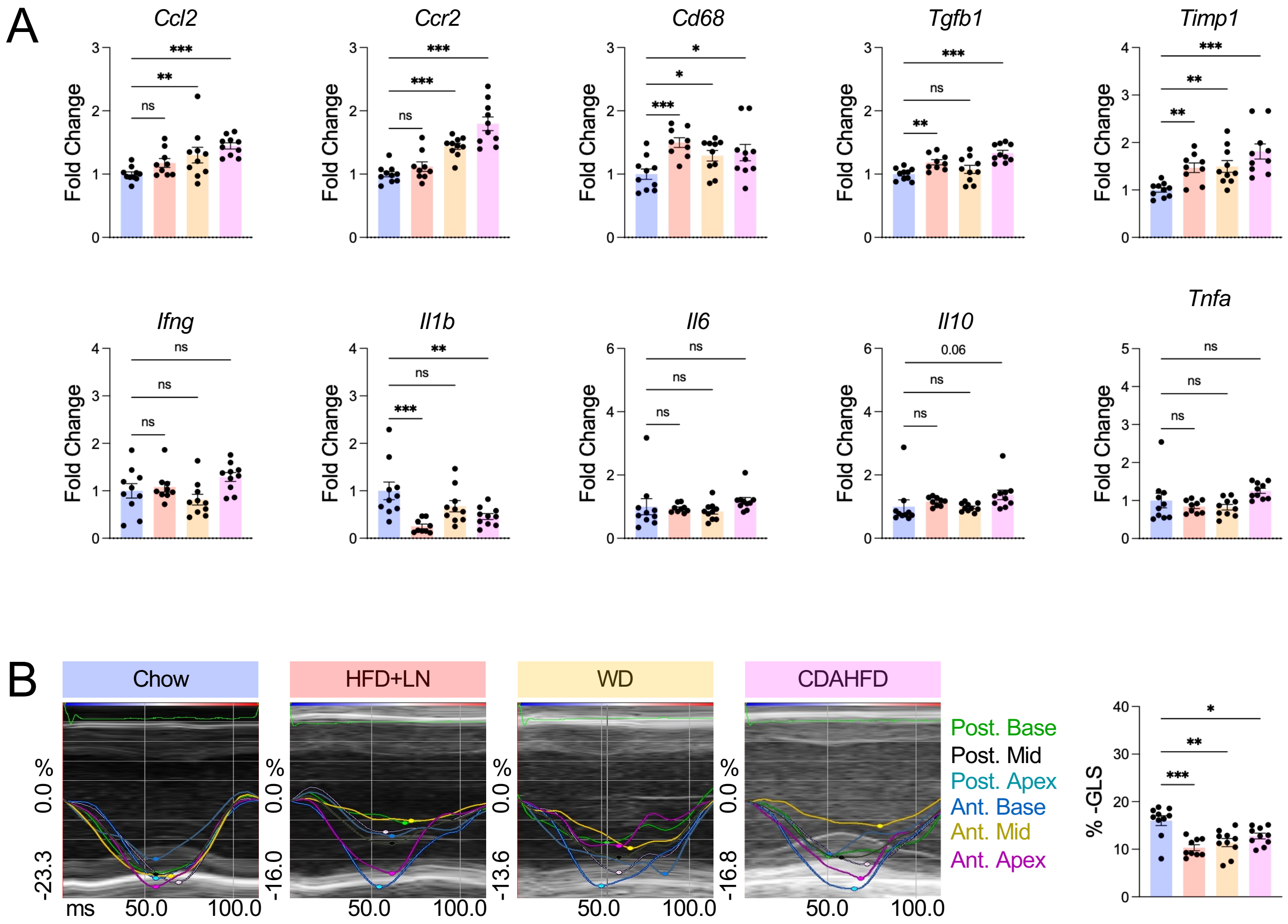


**Supplemental Figure 3. Cardiac gene expression and global longitudinal strain differ across complementary cardiometabolic stress models.** Male C57BL/6 mice were treated for 10 weeks prior to tissue collection. **A** Cardiac expression of inflammatory and fibrotic genes. **B** Global longitudinal strain (GLS) measured by speckle-tracking echocardiography. Data are presented as mean ± SEM. $n$ = 9–10 mice per group pooled from two or more independent experiments. *$P < 0.05$, **$P < 0.01$, ***$P < 0.001$ by one-way ANOVA followed by Tukey's multiple comparisons test.

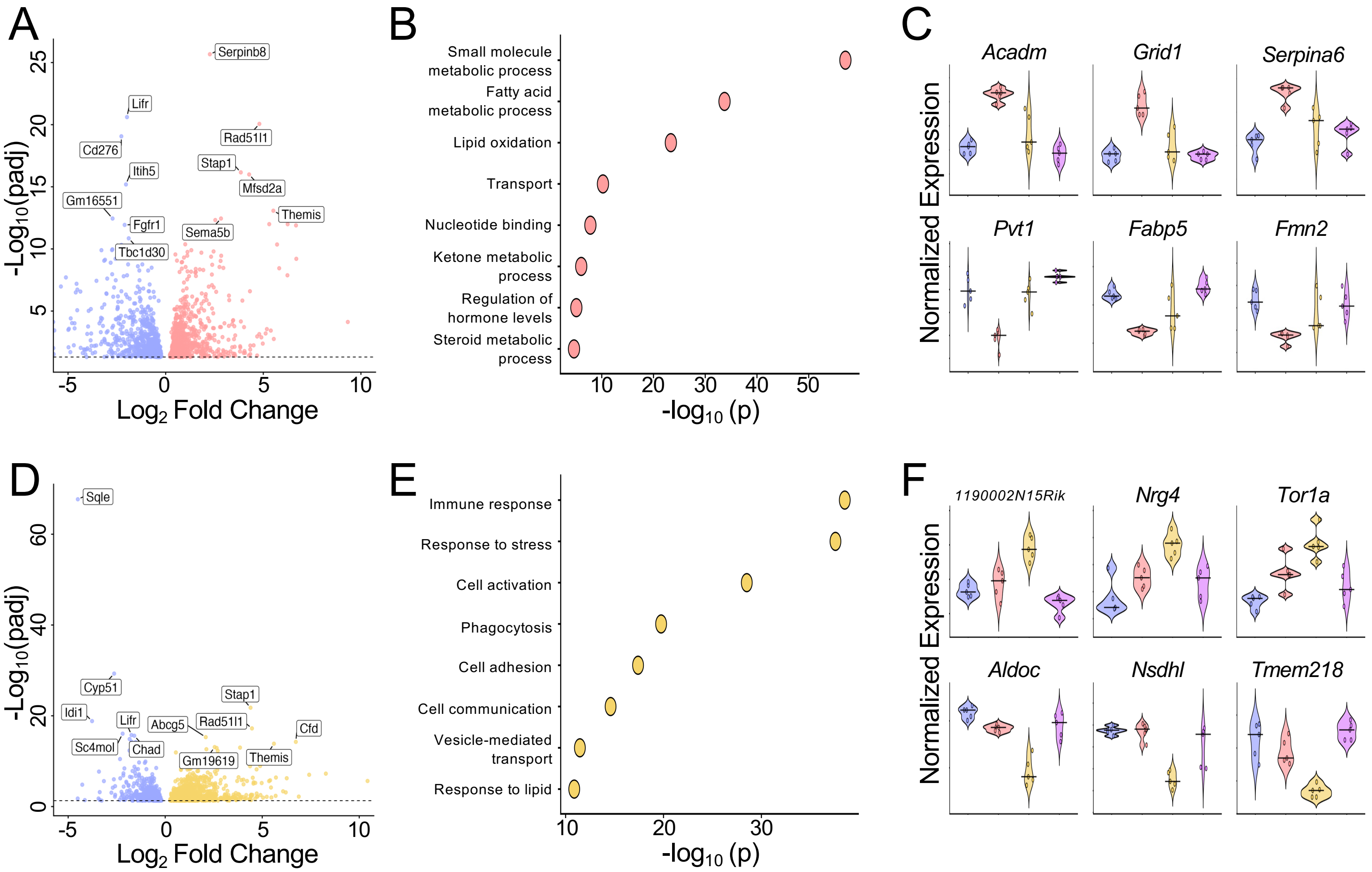


**Supplemental Figure 4. HFpEF and WD induce distinct hepatic transcriptomic responses.** Male C57BL/6 mice were treated for 10 weeks prior to liver tissue collection for bulk RNA sequencing. **A** Volcano plot of differential gene expression in HFD+LN versus chow. **B** Pathway enrichment analysis of differentially expressed genes upregulated in HFD+LN versus chow. **C** Expression of the three most upregulated and downregulated genes in HFD+LN compared with all experimental groups. **D** Volcano plot of differential gene expression in WD versus chow. **E** Pathway enrichment analysis of differentially expressed genes upregulated in WD versus chow. **F** Expression of the three most upregulated and downregulated genes in WD compared with all experimental groups. *n* = 5 mice per group pooled from two independent experiments.

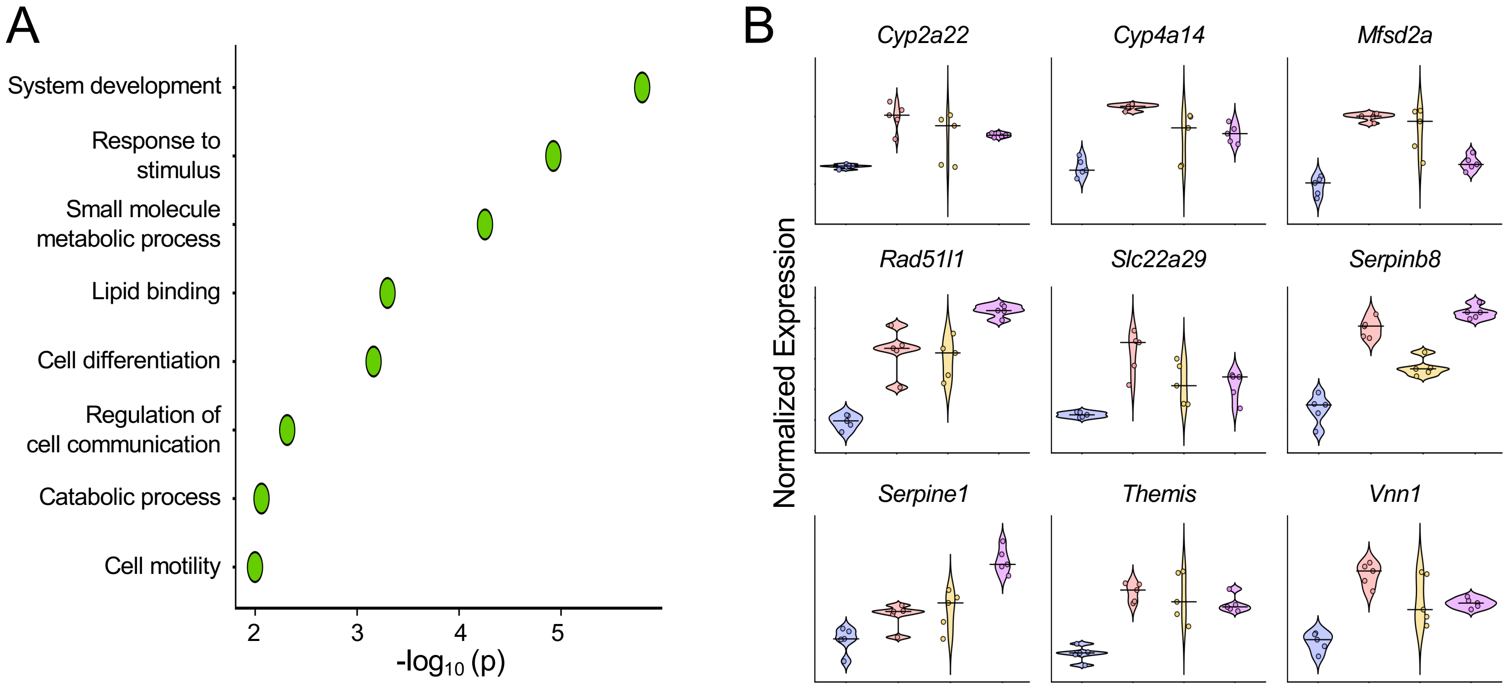


**Supplemental Figure 5. Integrated liver transcriptomics identifies conserved pathways and genes across complementary cardiometabolic stress models.** Male C57BL/6 mice were treated for 10 weeks prior to liver tissue collection for bulk RNA sequencing. Differentially expressed genes from HFD+LN, WD, and CDAHFD groups were identified relative to chow controls and integrated to define conserved hepatic responses across cardiometabolic stress models. **A** Pathway enrichment analysis of genes differentially increased across all three cardiometabolic disease conditions relative to chow, identifying conserved hepatic biological pathways. **B** Violin plots showing the normalized expression of conserved upregulated genes across experimental groups. $n$ = 5 mice/group pooled from two independent experiments.

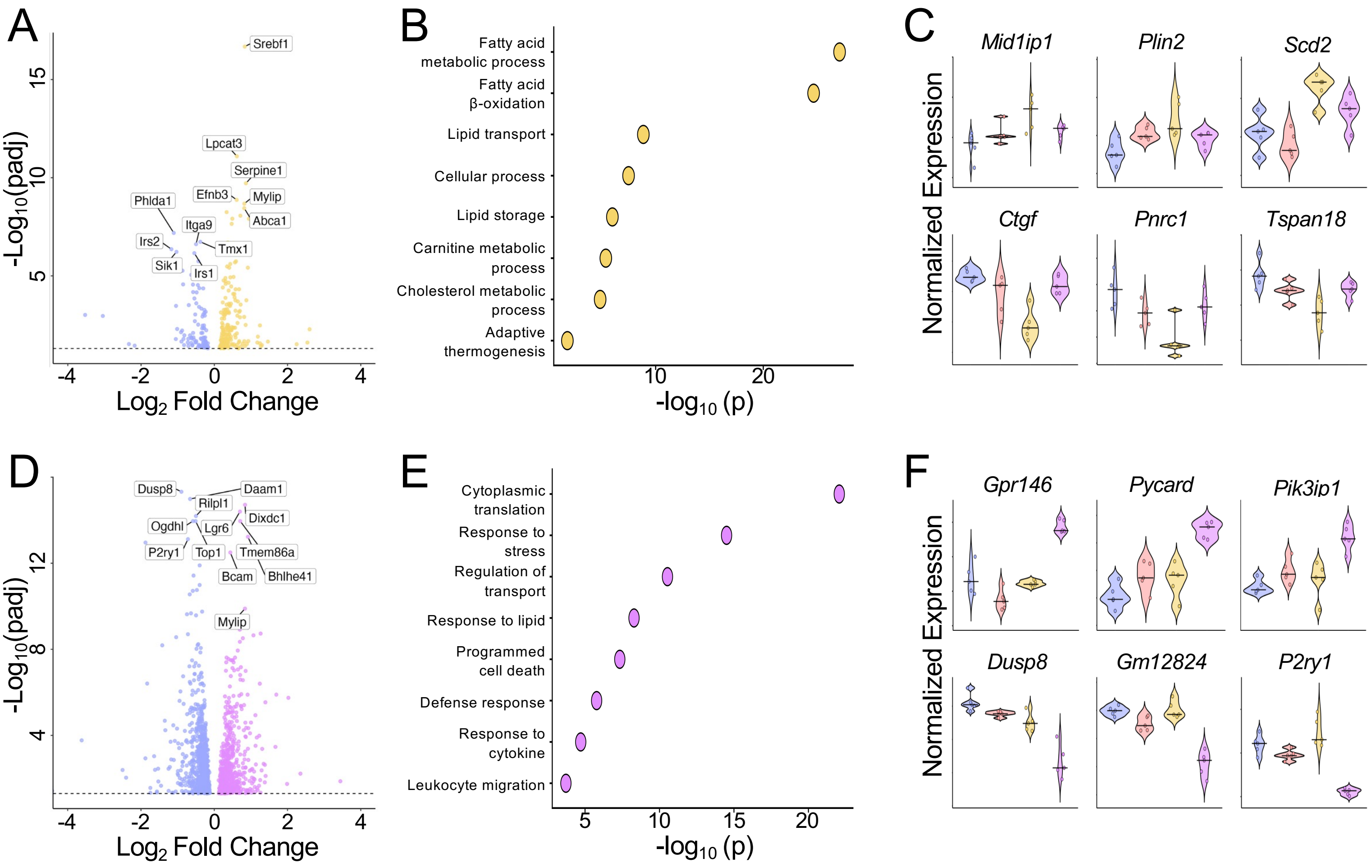


**Supplemental Figure 6. WD and CDAHFD induce distinct cardiac transcriptomic responses.** Male C57BL/6 mice were treated for 10 weeks prior to heart tissue collection for bulk RNA sequencing. **A** Volcano plot of differential gene expression in WD versus chow. **B** Pathway enrichment analysis of differentially expressed genes upregulated in WD versus chow. **C** Expression of the three most upregulated and downregulated genes in WD compared with all experimental groups. **D** Volcano plot of differential gene expression in CDAHFD versus chow. **E** Pathway enrichment analysis of differentially expressed genes upregulated in CDAHFD versus chow. **F** Expression of the three most upregulated and downregulated genes in CDAHFD compared with all experimental groups. *n* = 5 mice/group pooled from two independent experiments.

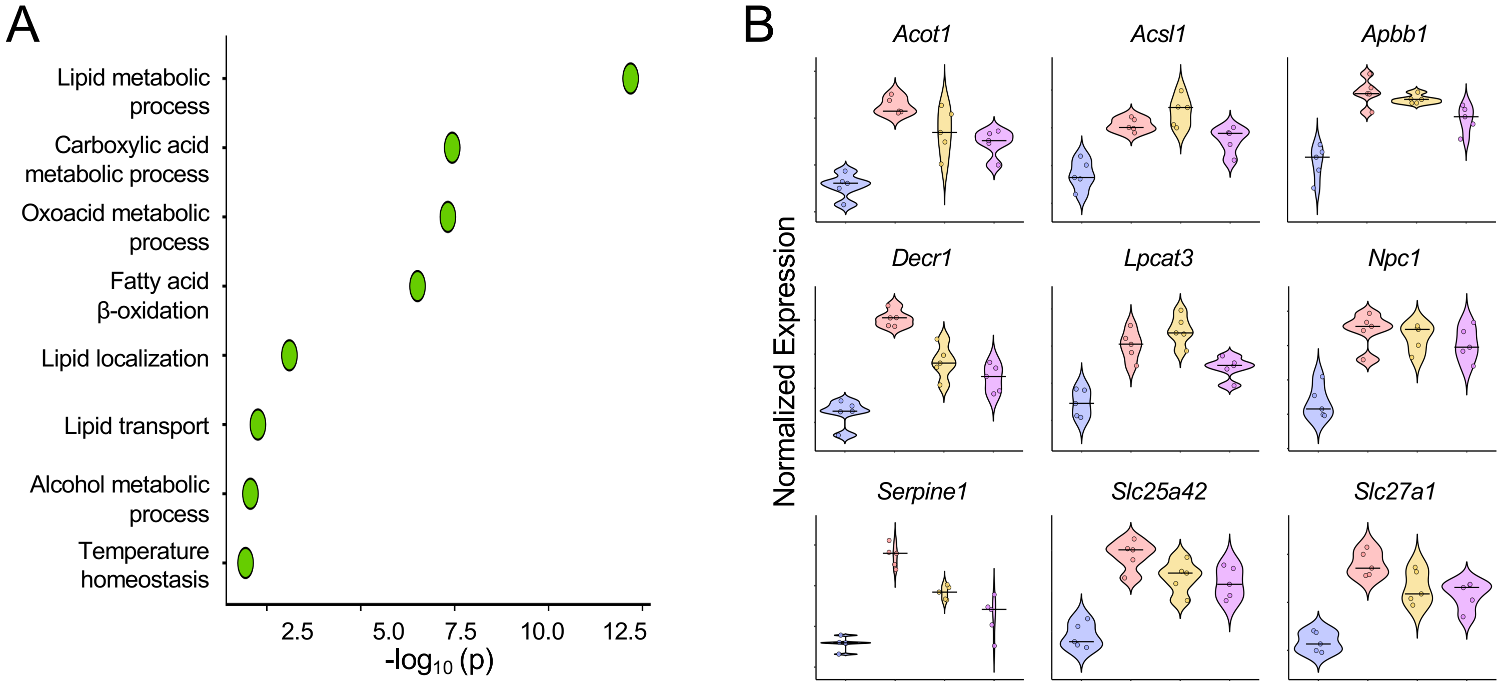


**Supplemental Figure 7. Integrated cardiac transcriptomics identifies conserved pathways and genes across complementary cardiometabolic stress models.** Male C57BL/6 mice were treated for 10 weeks prior to heart tissue collection for bulk RNA sequencing. Differentially expressed genes from HFD+LN, WD, and CDAHFD groups were identified relative to chow controls and integrated to define conserved cardiac responses across cardiometabolic stress models. **A** Pathway enrichment analysis of genes differentially increased across all three cardiometabolic disease conditions relative to chow. **B** Violin plots showing the expression of conserved upregulated genes across experimental groups. *n* = 5 mice/group pooled from two independent experiments.
